## Supplementary Information for "Apoptotic signaling clears engineered *Salmonella* in an organ-specific manner"

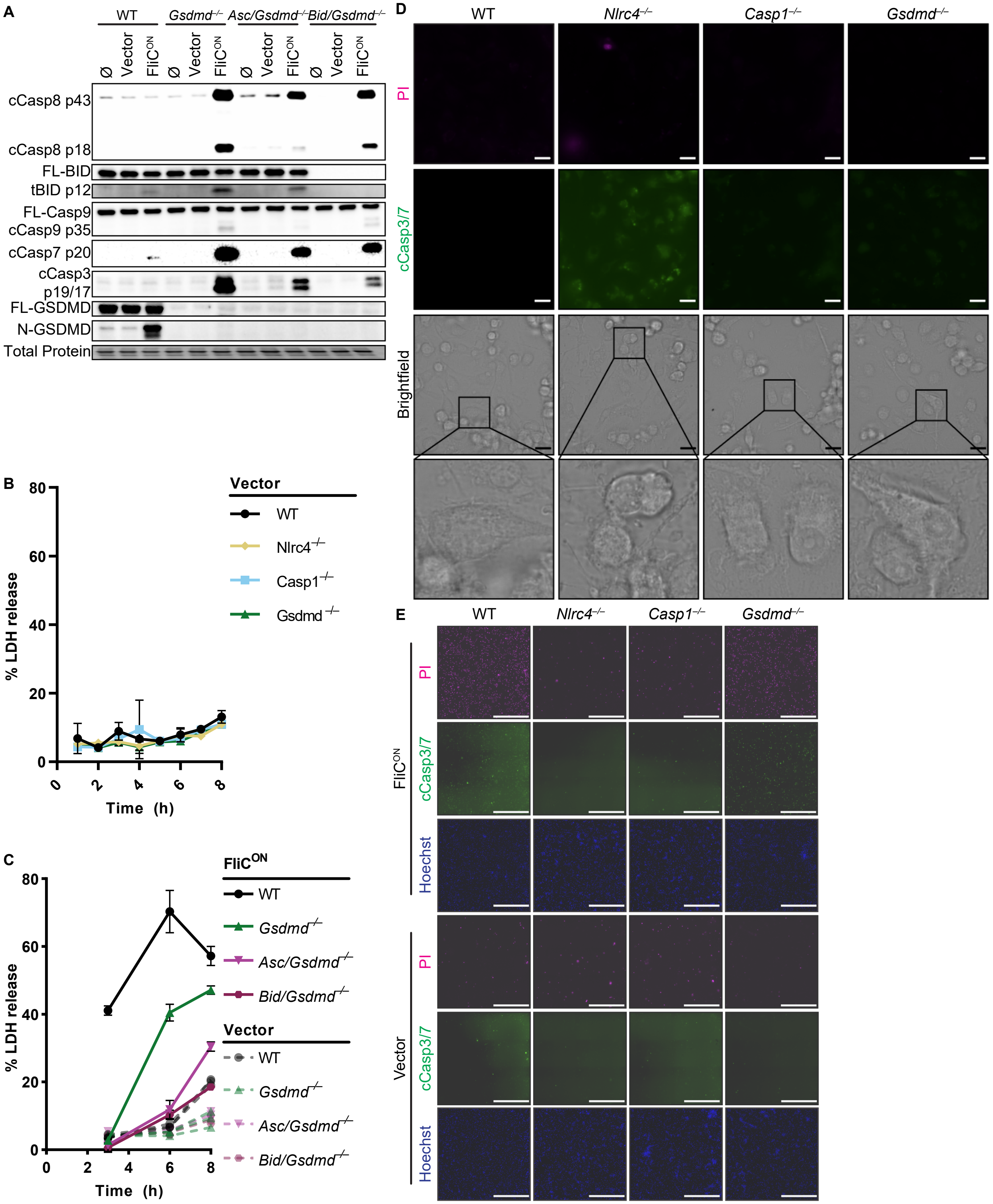

A
WT
Gsdmd−/−
Asc/Gsdmd−/−
Bid/Gsdmd−/−
Ø
Vector
FliC^ON
cCasp8 p43
cCasp8 p18
FL-BID
tBID p12
FL-Casp9
cCasp9 p35
cCasp7 p20
cCasp3
p19/17
FL-GSDMD
N-GSDMD
Total Protein
B
% LDH release
Time (h)
Vector
WT
Nlrc4−/−
Casp1−/−
Gsdmd−/−
C
% LDH release
Time (h)
FliC^ON
WT
Gsdmd−/−
Asc/Gsdmd−/−
Bid/Gsdmd−/−
Vector
WT
Gsdmd−/−
Asc/Gsdmd−/−
Bid/Gsdmd−/−
D
WT
Nlrc4−/−
Casp1−/−
Gsdmd−/−
PI
cCasp3/7
Brightfield
E
WT
Nlrc4−/−
Casp1−/−
Gsdmd−/−
FliC^ON
PI
cCasp3/7
Hoechst
Vector
PI
cCasp3/7
Hoechst

**Fig. S1. Vector control *S.* Typhimurium does not activate apoptotic backup pathways in vitro.**

A-E) BMMs were infected with indicated SPI2-induced *S*. Typhimurium strains.

A) Western blot analysis of whole cell lysates at 4 hpi. Results from one experiment.

B) LDH release at 1-8 hpi. Results representative of 3 independent experiments. Data are represented as mean ± SD of 3 technical replicates. Data performed at the same time as Figure 1E, graphed separately for visualization.

C) LDH release at 1-8 hpi. Results from one experiment. Data are represented as mean ± SD of 3 technical replicates.

D-E) Immunofluorescence and brightfield at 4hpi. Cells were stained with PI, cleaved caspase-3/7, and Hoechst. Representative image from two (brightfield, PI) or one (cleaved caspase-3/7) independent experiments. Data performed at the same time as Figure 1F. D) 60x magnification, scale bar 20 μm. E) 20x stitched image, scale bar 500 μm.

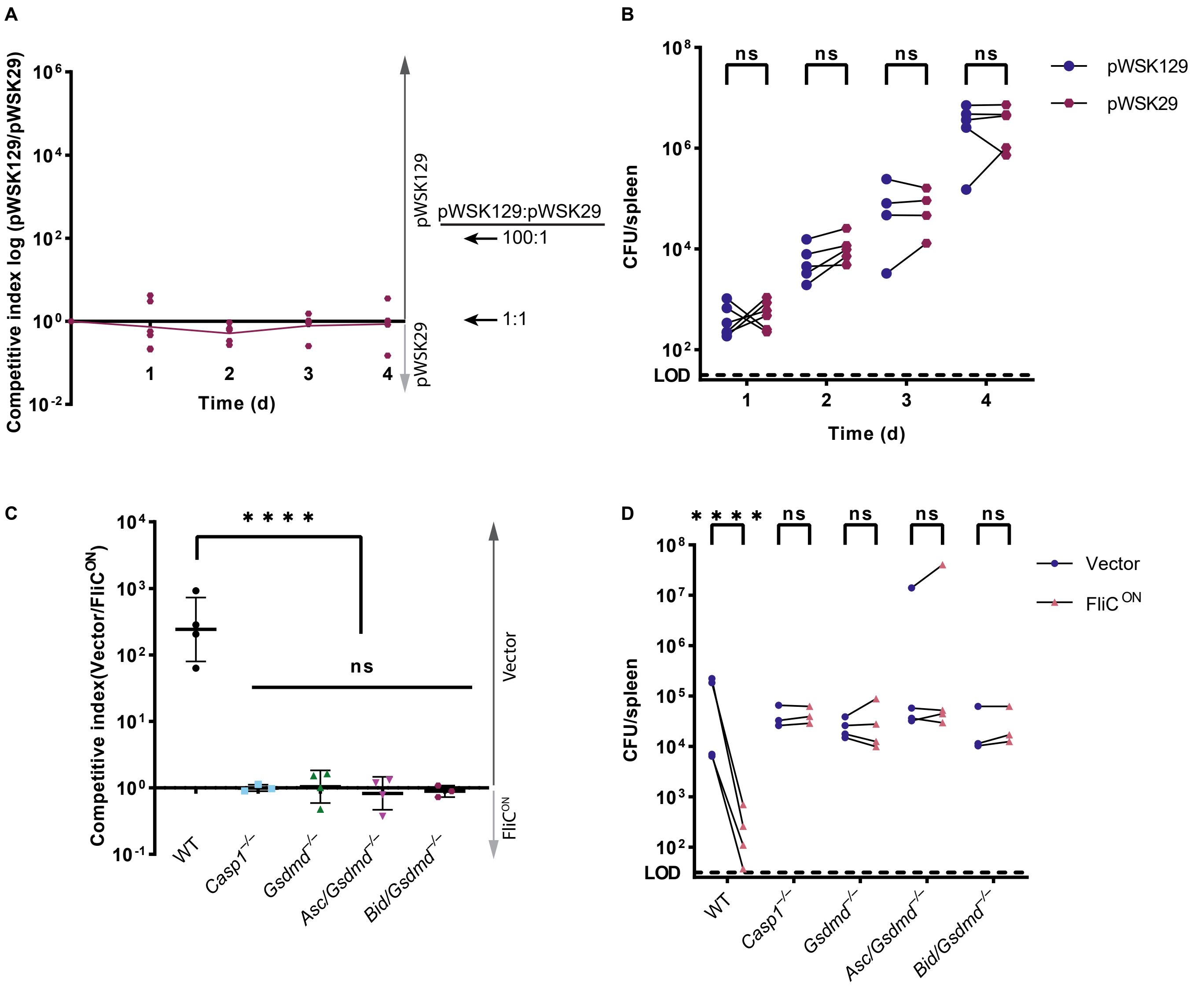

A
Competitive index log (pWSK129/pWSK29)
10^6
10^4
10^2
10^0
10^-2
1
2
3
4
Time (d)
pWSK129
pWSK29
pWSK129:pWSK29
100:1
1:1
B
10^8
10^6
10^4
10^2
LOD
ns
ns
ns
ns
pWSK129
pWSK29
CFU/spleen
1
2
3
4
Time (d)
C
Competitive index(Vector/FliC^ON)
10^4
10^3
10^2
10^1
10^0
10^-1
* * * *
ns
Vector
FliC^ON
WT
Casp1^-/-
Gsdmd^-/-
Asc/Gsdmd^-/-
Bid/Gsdmd^-/-
D
* * * *
ns
ns
ns
ns
10^8
10^7
10^6
10^5
10^4
10^3
10^2
LOD
CFU/spleen
Vector
FliC ON
WT
Casp1^-/-
Gsdmd^-/-
Asc/Gsdmd^-/-
Bid/Gsdmd^-/-

**Fig. S2. Competitive index model can be used to study clearance of *S.* Typhimurium in vivo.**

A-B) Mice were infected with a 1:1 ratio of pWSK129 (“vector”) and pWSK29 (backbone of FliC$^{ON}$ and BID$^{ON}$ plasmids) *S*. Typhimurium. Mice were infected with $5x10^2$ CFU of each strain. Bacterial burdens in the spleen were determined at the indicated timepoints.

A) Timecourse competitive index infection in WT mice. Ratio of vector to pWSK29 is graphed. Data from one independent experiment, line connects means, n=4-6 mice per timepoint.

B) Individual burdens of vector and pWSK29 from (A). Paired vector and pWSK29 data from each mouse are connected by a line. Two-way ANOVA n.s. $p<0.05$

C-D) Mice were infected with a 1:1 ratio of FliC$^{ON}$ and a vector control *S*. Typhimurium. Mice were infected with $5x10^2$ CFU of each strain. Bacterial burdens in the spleen were determined at 48 hpi.

C) Competitive index infection in indicated mice. Data from one independent experiment, line representing mean ± SD, n=3-4 mice per genotype. One-way ANOVA n.s. $p>0.05$; ****$p<0.0001$.

D) Individual burdens of vector and FliC$^{ON}$ from (C). Paired vector and FliC$^{ON}$ data from each mouse are connected by a line. Two-way repeated measure ANOVA. n.s. $p>0.05$, ****$p<0.0001$

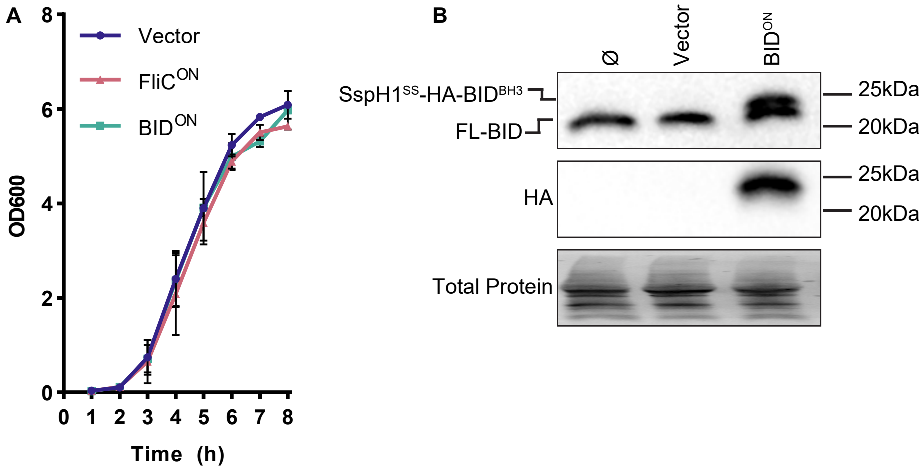

A
B
Vector
FliC^ON
BID^ON
OD600
Time (h)
Ø
Vector
BID^ON
SspH1^SS-HA-BID^BH3
FL-BID
25kDa
20kDa
HA
Total Protein

**Fig S3. Production of SspH1$^{SS}$-HA-BID$^{BH3}$ construct does not cause growth defects in BID$^{ON}$ *S*. Typhimurium.**
A) OD600 growth curve in LB media.
B) BMMs were infected with SPI2-induced *S.* Typhimurium. Western blot analysis of whole cell lysates at 6 hpi. Double band of endogenous full length BID and sspH1$^{SS}$-HA-BID$^{BH3}$ resolved.

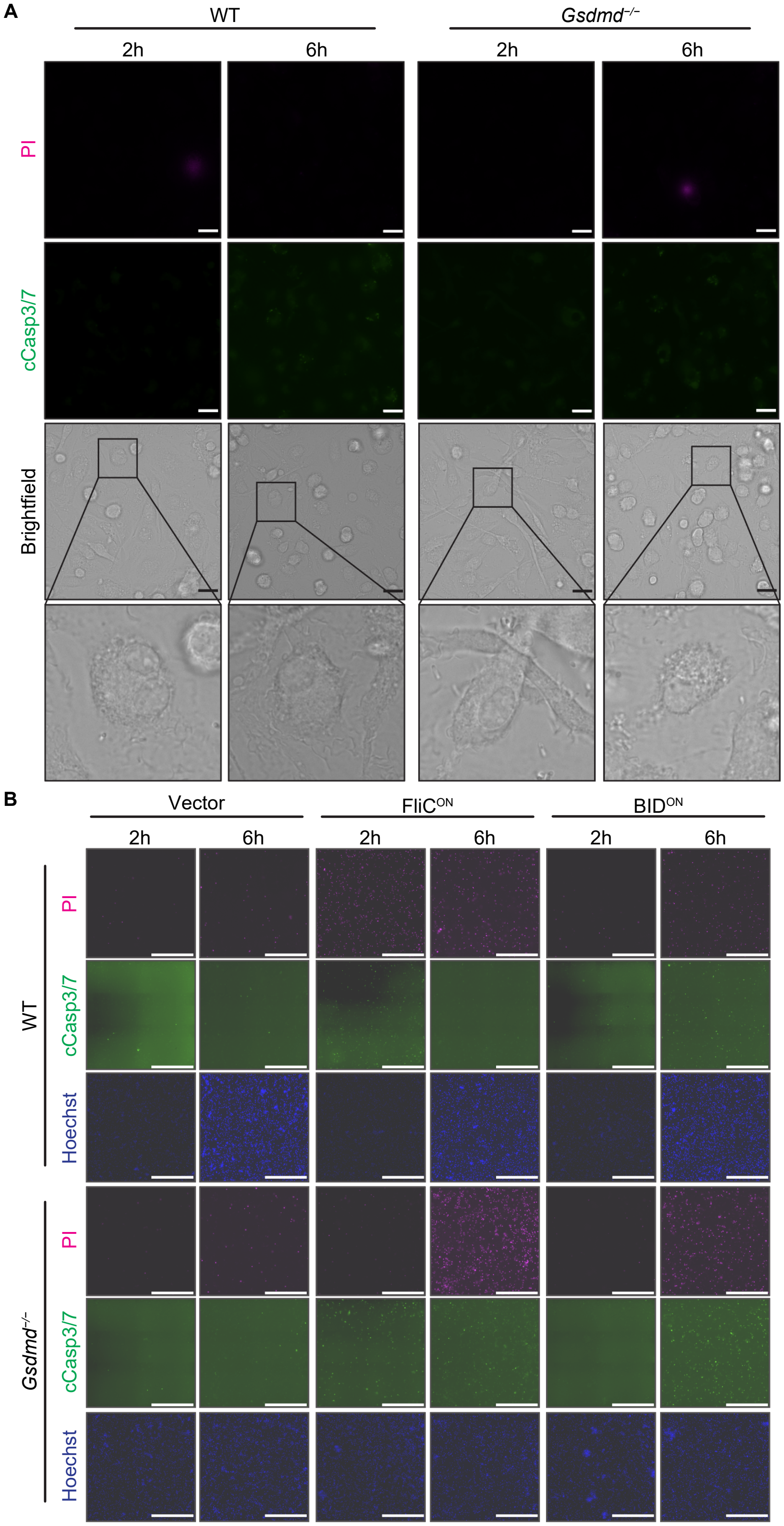
A
WT
Gsdmd−/−
2h
6h
2h
6h
PI
cCasp3/7
Brightfield
B
Vector
FliC^ON
BID^ON
2h
6h
2h
6h
2h
6h
WT
PI
cCasp3/7
Hoechst
Gsdmd−/−
PI
cCasp3/7
Hoechst

**Fig S4. Vector control *S*. Typhimurium does not cause RCD in vitro.**

A-B) BMMs were infected with indicated SPI2-induced *S*. Typhimurium strains. Cells were stained with PI, cleaved caspase-3/7, and Hoechst and imaged at indicated timepoints. Representative image from three (brightfield, PI) or one (cleaved caspase-3/7) independent experiments. Data performed at the same time as Figure 4B. A) 60x magnification, scale bar 20 μm. B) 20x stitched image, scale bar 500 μm.

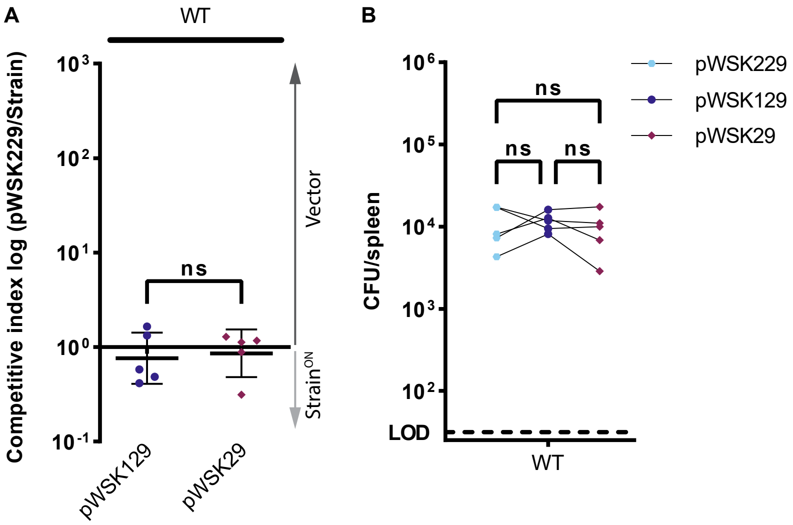

A
WT
Competitive index log (pWSK229/Strain)
10^3
10^2
10^1
10^0
10^-1
ns
Vector
Strain^ON
pWSK129
pWSK29
B
CFU/spleen
10^6
10^5
10^4
10^3
10^2
LOD
ns
ns
ns
WT
pWSK229
pWSK129
pWSK29

**Fig S5. pWSK229 can be used for triple competitive index infection in vivo.**
A-B) Mice were infected simultaneously with three strains, $5 \times 10^2$ CFU each of pWSK229 ("vector (Cam)", Cam), pWSK129 (Kan), and pWSK29 (Amp) *S*. Typhimurium. Bacterial burdens in the spleen were determined at 48 hpi.
A) Triple competitive index infection of WT mice. Ratio of vector (Cam) to pWSK129 and vector to pWSK29 is graphed. Data representative of three experiments. Data represented by line at mean ± SD, n=5. Unpaired two-tailed t-test n.s. $p>0.05$.
B) Individual burdens of vector (Cam), pWSK129, and pWSK29 from (A). Paired vector, pWSK129, and pWSK29 data from each mouse are connected by a line. Repeated measures one-way ANOVA n.s. $p>0.05$.

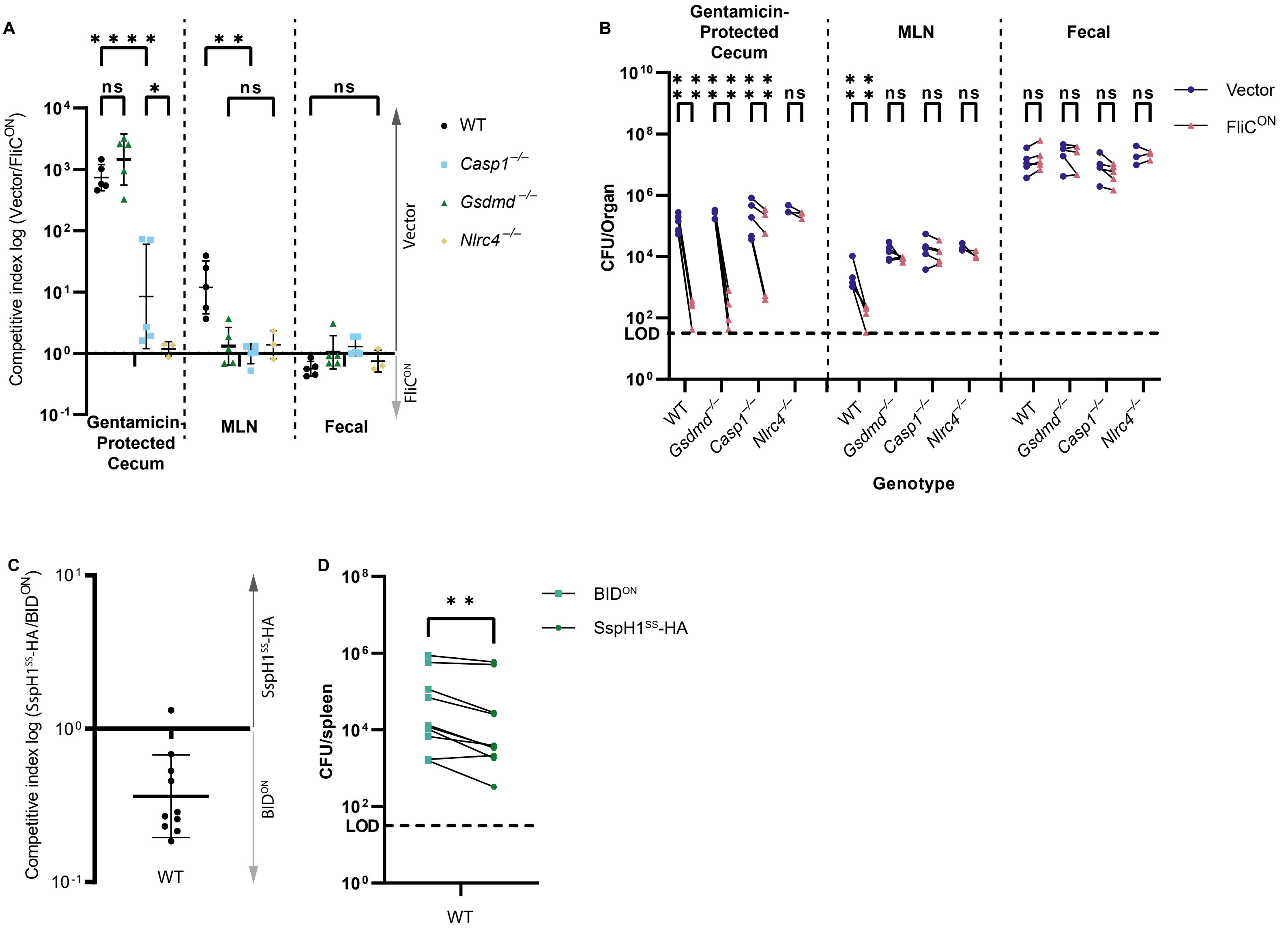

A
Competitive index log (Vector/FliC^ON)
10^4
10^3
10^2
10^1
10^0
10^-1
****
ns
*
**
ns
ns
Gentamicin-Protected Cecum
MLN
Fecal
Vector
FliC^ON
WT
Casp1−/−
Gsdmd−/−
Nlrc4−/−
B
Gentamicin-Protected Cecum
MLN
Fecal
CFU/Organ
10^10
10^8
10^6
10^4
10^2
10^0
LOD
****
****
****
ns
**
Vector
FliC^ON
WT
Gsdmd−/−
Casp1−/−
Nlrc4−/−
Genotype
C
Competitive index log (SspH1^SS-HA/BID^ON)
10^1
10^0
10^-1
SspH1^SS-HA
BID^ON
WT
D
CFU/spleen
10^8
10^6
10^4
10^2
10^0
LOD
**
BID^ON
SspH1^SS-HA
WT

**Fig S6. Clearance of $FliC^{ON}$ in the cecum is NLRC4-dependent**

A-B) Mice were orally treated with 20mg streptomycin, and 24h later orally infected with $1x10^7$ CFUs total bacteria comprised of a 1:1 ratio of $FliC^{ON}$ and vector control *S.* Typhimurium, all on a *flgB* mutant background. Bacterial burdens in the cecum, mesenteric lymph nodes (MLN), and fecal sample were determined at 48 hpi.

A) Competitive index is graphed as a ratio of vector to $FliC^{ON}$. Data from one independent experiment, line representing mean ± SD, n=3-6 mice per condition. Two-way repeated measure ANOVA n.s. p>0.05, *p<0.05, **p<0.01, ****p<0.0001.

B) Individual burdens of vector and $FliC^{ON}$ from (A). Paired vector and $FliC^{ON}$ data from each mouse are connected by a line. Two-way repeated measure ANOVA n.s. p>0.05, ****p<0.0001.

C-D) Mice were infected with a 1:1 ratio of $Kan^R$ $BID^{ON}$ and $SspH1^{SS}$-HA vector control *S.* Typhimurium. Mice were infected with $5x10^2$ CFU of each strain. Bacterial burdens in the spleen were determined at 48 hpi.

C) Competitive index infection of WT mice infected with $Kan^R$ $BID^{ON}$. Ratio of $SspH1^{SS}$-HA vector to $Kan^R$ $BID^{ON}$ is graphed. Data is combined from two independent experiments, line representing mean ± SD, n=10 mice.

D) Individual burdens of $SspH1^{SS}$-HA vector and $Kan^R$ $BID^{ON}$ from (C). Paired $SspH1^{SS}$-HA vector and $Kan^R$ $BID^{ON}$ data from each mouse are connected by a line. Paired t-test **p<0.01

**Movie S1 (separate file). Engineered FliC$^{ON}$ *S*. Typhimurium activates apoptosis in *Gsdmd*$^{-/-}$ BMMs in vitro.**

*Gsdmd*$^{-/-}$ BMMs were infected with FliC$^{ON}$ SPI2-induced *S.* Typhimurium strains. Cells were stained with PI, cleaved caspase-3/7, and Hoechst and imaged at 6 hpi. 60x magnification, scale bar 20µm. Z-stack of data in Figure 1G (slices 11, 18, 21, 26) and Figure 4B (slice 19). Performed in the same experiment as Movie S2.

**Movie S2 (separate file). Engineered BID$^{ON}$ *S*. Typhimurium causes apoptosis in WT BMMs in vitro.**

WT BMMs were infected with BID$^{ON}$ SPI2-induced *S.* Typhimurium strains. Cells were stained with PI, cleaved caspase-3/7, and Hoechst and imaged at 6 hpi. 60x magnification, scale bar 20µm. Z-stack of data in Figure 3H (slices 11, 19, 22, 24) and Figure 4B (slice 20). Performed in the same experiment as Movie S1.
